## Supplementary figures and images for "Multi-omics investigation of spontaneous T2DM macaque emphasizes gut microbiota could up-regulate the absorption of excess palmitic acid in the T2DM progression"

### Figure S1

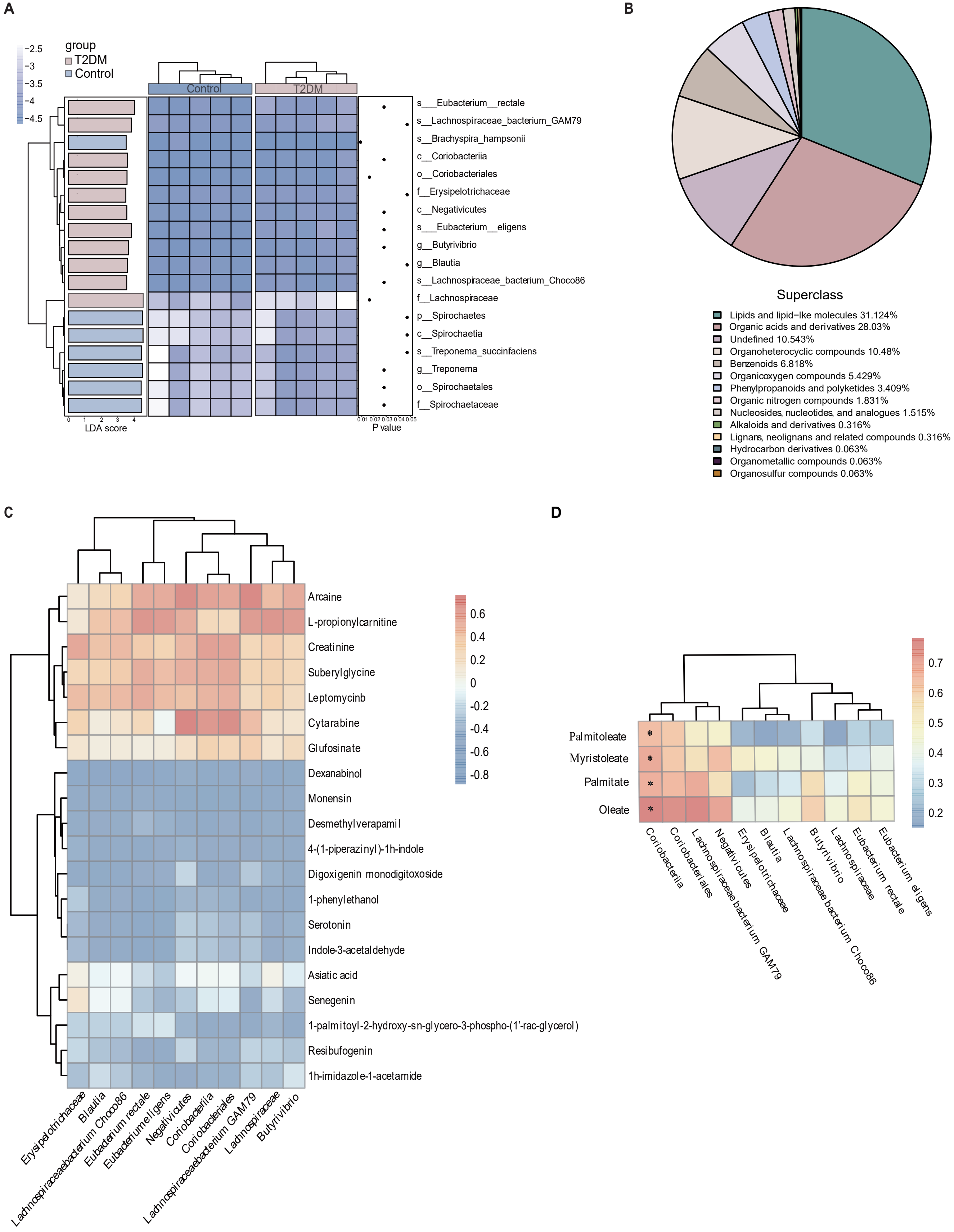

### Figure S2

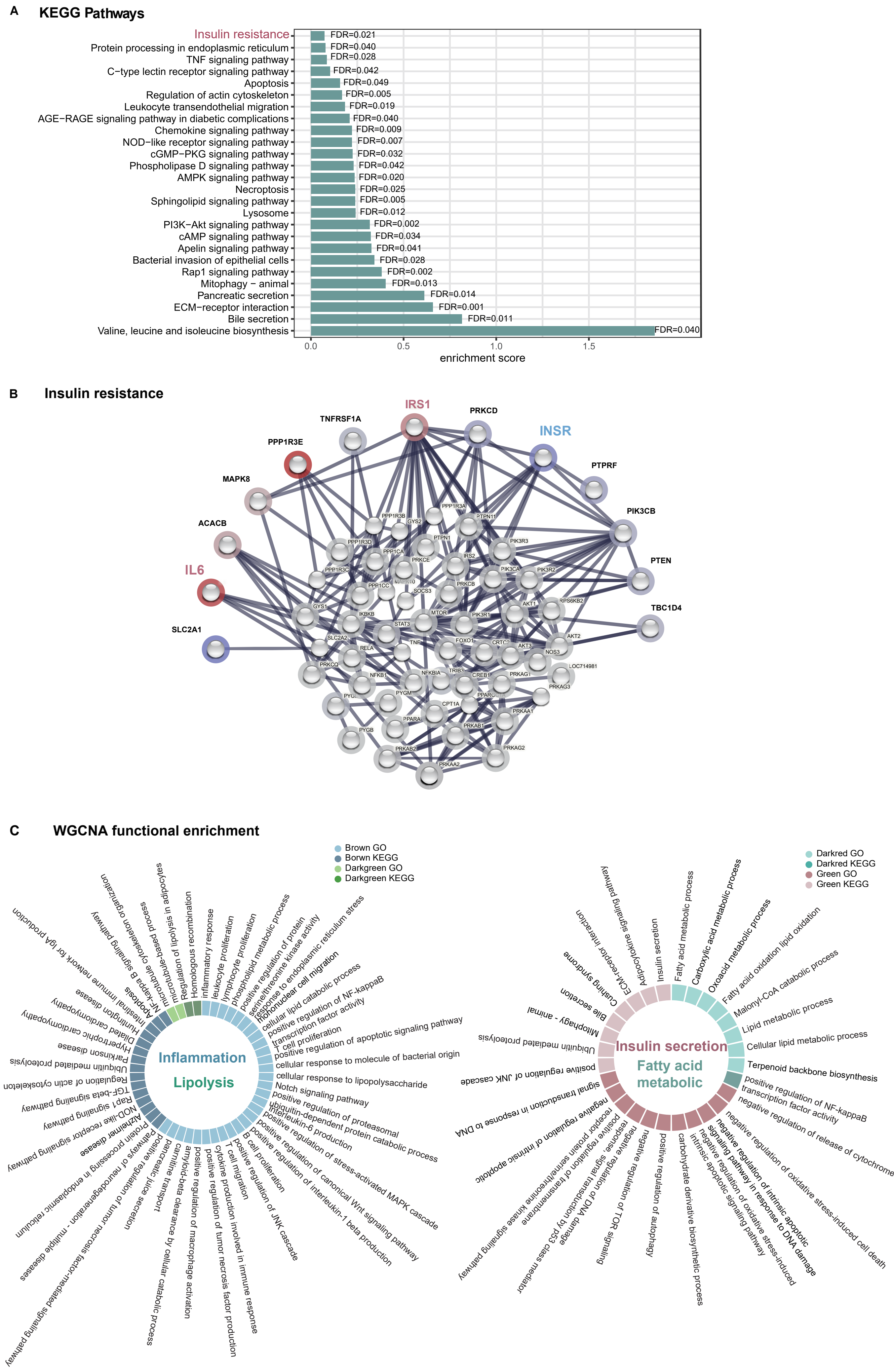

### Figure S3

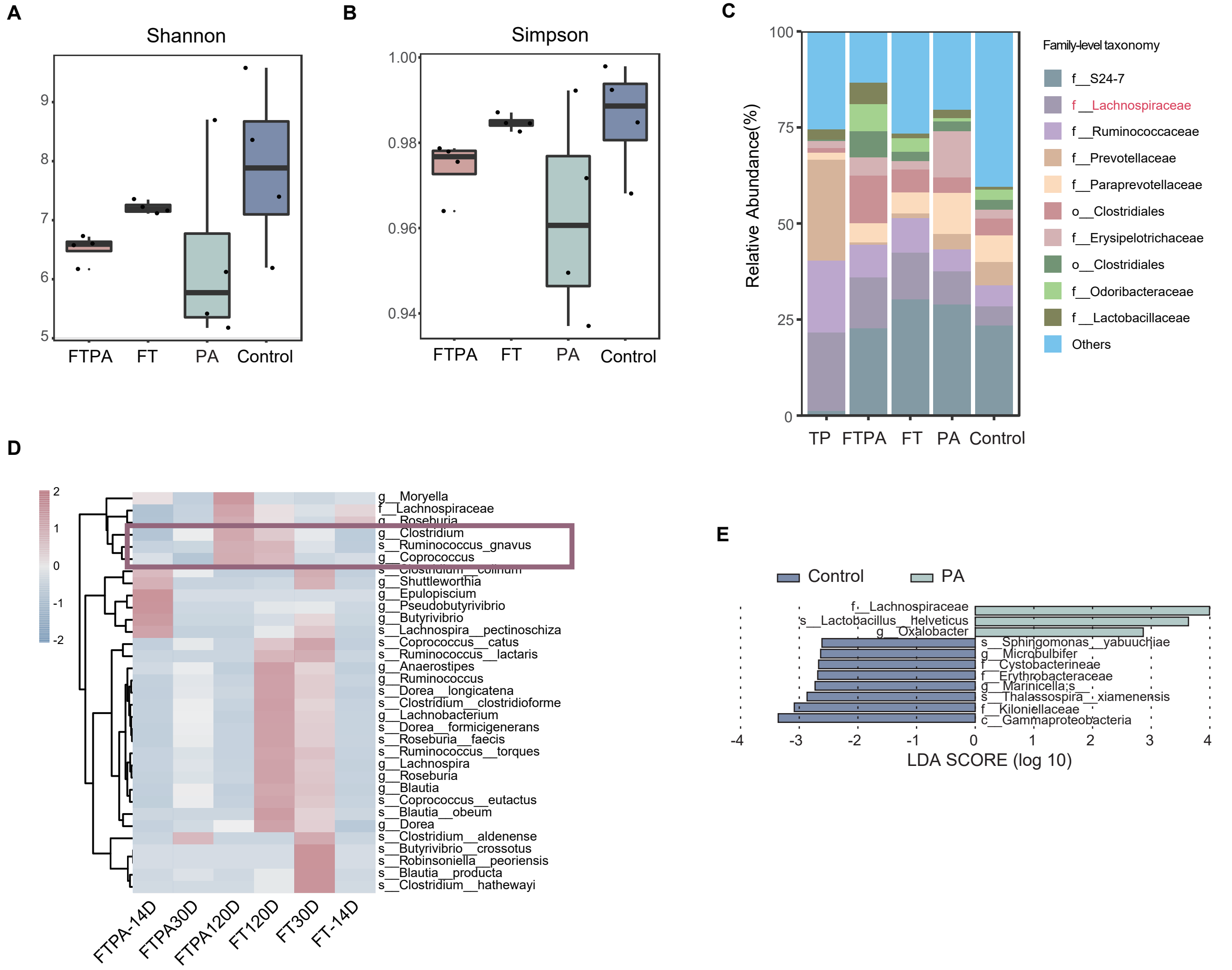

### Figure S4

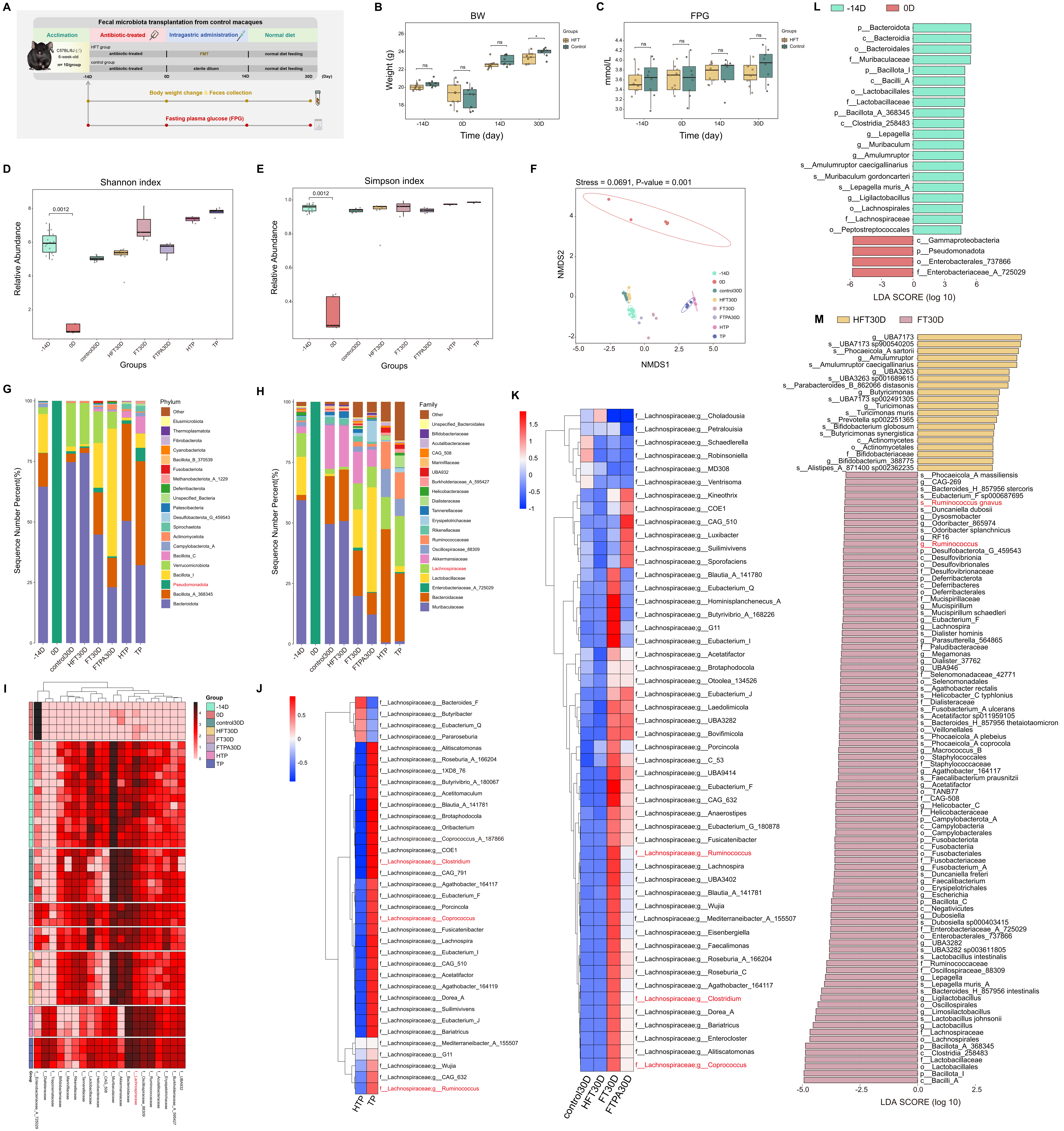
